# GATA4, expressed in Barrett’s esophagus and esophageal adenocarcinoma, can block squamous epithelial cell gene expression in human esophageal cells

**DOI:** 10.1101/2020.06.17.156026

**Authors:** Roman Stavniichuk, Ann DeLaForest, Cayla A. Thompson, James Miller, Rhonda F. Souza, Michele A. Battle

## Abstract

Metaplasia often involves a change from one cell type to another that was present during organogenesis. The embryonic esophagus is initially lined by columnar cells that are replaced by squamous cells, and metaplasia in Barrett’s esophagus (BE) involves a change from squamous to columnar cells in the setting of gastroesophageal reflux. Here, we explored the effect of ectopic expression of the essential developmental transcription factor GATA4 on squamous epithelial cell gene expression using human esophageal squamous epithelial cells. We found that GATA4 protein, although absent in mature human esophageal squamous epithelium, was present in BE and esophageal adenocarcinoma (EAC). Moreover, acid and bile induced *GATA4* mRNA in esophageal squamous epithelial cells. Ectopic GATA4 expression in esophageal squamous epithelial cells generally compromised squamous cell marker gene expression, although the extent varied between cell lines studied. We observed GATA4 occupancy in the *p63, KRT5*, and *KRT15* gene promoters, suggesting that GATA4 can directly repress expression of typical squamous epithelial cell marker genes. Overall, our data suggest a mechanism whereby GATA4 expression in abnormal esophageal cells, possibly induced by reflux, supports a columnar metaplastic cell identity by repressing expression of key genes required to program stratified squamous epithelial cell identity.

## INTRODUCTION

Gastroesophageal reflux disease (GERD) is the most prevalent gastrointestinal disorder in the US with as many as 20% of the population experiencing weekly GERD symptoms.[**1**] A serious consequence of chronic GERD and its associated inflammatory and cellular stress is development of Barrett’s esophagus (BE), a condition in which a metaplastic columnar epithelium with gastric and intestinal features replaces the normal stratified squamous esophageal epithelium.[**2–7**] BE, estimated to affect 5.6% of American adults, is most alarming because it is a significant risk factor for esophageal adenocarcinoma (EAC), an aggressive and deadly cancer with a five-year survival rate of only 19% that has increased in incidence by more than 400% since 1975.[**3**,**4**,**8**,**9**] Esophageal cancer is most often diagnosed after symptoms have appeared, marking an advanced stage of the disease.[**10**] Given esophageal cancer’s aggressive nature and poor prognosis along with a lack of non-invasive, early screening methods, the mechanisms controlling the progression of normal tissues to BE to EAC in GERD patients has been the subject of intense scrutiny. Although it is well established that chronic GERD and associated reflux-induced inflammation and cellular damage drive development of BE, the precise molecular events driving metaplasia are not well understood. Several mechanisms to explain BE have been proposed, but none have yet been definitively proven in human disease.[**4–7**] One possibility is that reflux-induced tissue damage and inflammation cause cellular reprogramming of esophageal stem/progenitor cells.[**11–14**] Cells residing in submucosal esophageal glands have also been identified as a possible BE origin.[**15**] Alternatively, stem cells from the stomach or from the bone marrow have also been implicated in BE.[**16–19**] Most recently, a population of unique transitional cells residing at the junction between the stratified squamous epithelium of the esophagus and simple columnar epithelium of the stomach has been identified as a potential source of BE cells.[**20**] Importantly, these mechanisms are not mutually exclusive, and BE may emerge via multiple routes involving resident esophageal cells and cells from other tissue origins including the transitional and cardiac zones. Regardless of the origin of metaplastic BE cells, however, changes in transcriptional activity are paramount in BE pathogenesis. For example, changes in transcription factor activity would be necessary to re-program esophageal stem cells to generate the metaplastic columnar epithelium characteristic of BE or to direct migration of cells from other tissues into the damaged esophageal region. Altered transcriptional programs likely also contribute to the maintenance of abnormal, metaplastic columnar epithelium in the esophagus. Therefore, transcription factors with altered expression and activity in BE compared with normal tissue are primary candidates to evaluate in BE pathogenesis. Defining such factors and their downstream targets may lead to identification of novel BE therapeutic targets and/or biomarkers.

Several columnar cell type transcription factors, expressed in human BE and EAC, have been shown to drive varying degrees of metaplasia in esophageal cells or tissue. When expressed in mouse three-dimensional organotypic esophageal cultures, SOX9 induces expression of columnar-type cytokeratins and converts the multilayer structure to a 1-2 cell thick structure with cuboidal/columnar cell morphology.[**21**] Squamous lesions develop in the intestine of CDX2 heterozygous mice.[**22**] Squamous epithelial cells expressing CDX2 driven by the *KRT14* promoter are abnormal resembling multi-layer esophagus, a stage proposed as a precursor to full BE metaplasia.[**23**] Furthermore, CDX2 expression in transitional cells at the squamocolumnar junction using the *Krt7* promoter causes a BE-like metaplasia.[**20**] Co-expression of CDX2 and Myc in an immortalized esophageal squamous cell line, in conjunction with Notch inhibition, reduces squamous keratin expression and induced columnar keratin and mucin expression.[**24**] Finally, overexpression of HNF4A and FOXA2 in squamous cells also induces expression of columnar type transcripts and proteins including cytokeratins, villin, trefoil factor, and mucins.[**14**,**25**] On the other hand, loss of key squamous type transcription factors, including p63 and SOX2, has been shown to cause columnar metaplasia in mice. For example, embryos lacking p63 develop a BE-like metaplasia, and hypomorphic SOX2 mutants also develop regional columnar metaplasia in the esophagus and forestomach.[**26**,**27**]

In this study, we explored the effect of aberrant expression of GATA4 on the epithelial character of human esophageal squamous epithelial cells. GATA4 has been reported as expressed in human BE, and amplification of and expression of the *GATA4* gene has also been reported in EAC.[**28**,**29**] We hypothesized that ectopic expression of GATA4 in squamous esophageal epithelial cells in culture would alter the stratified squamous identity of the cells, possibly shifting these cells to a columnar, BE-like identity. We validated that GATA4 protein was robustly expressed in human BE and EAC but not in normal esophagus. We found that exposure of human esophageal squamous epithelial cells to acid and bile, two key reflux components implicated in BE etiology, induced *GATA4* mRNA expression. We expressed GATA4 protein in esophageal squamous epithelial cells and observed a loss of squamous epithelial cell marker expression at both mRNA and protein levels. We also demonstrated that GATA4 binds to consensus binding sites in the promoters of several squamous epithelial cell defining genes, including the master squamous epithelial cell transcriptional regulator p63, suggesting that GATA4 directly represses expression of squamous epithelial cell markers to suppress squamous cell fate in columnar tissues. Extrapolating to BE, the data suggest that GATA4 expression in BE cells would contribute to favoring a columnar cell identity rather than an esophageal squamous cell identity.

## RESULTS

### GATA4 is robustly expressed in human BE and EAC

BE is characterized by a squamous to columnar metaplasia, and as such, BE cells express genes typical of columnar epithelial cells. Given that GATA4 is a transcription factor expressed in columnar GI epithelial cells, we examined the extent to which BE and EAC tissues express GATA4. Two previous studies reported expression of GATA4 in human BE samples and amplification of the GATA4 gene in human EAC samples, but protein expression was not demonstrated in either study.[**28**,**29**] Therefore, we obtained biopsy samples from BE (n=10) and EAC (n=7) patients and stained for GATA4 protein. We detected GATA4 protein in metaplastic regions of all BE biopsies (Fig. 1). We further detected GATA4 protein in all EAC biopsies, although the intensity of the staining varied among and within tumors (Fig. 1). As expected, none of the normal esophageal biopsy samples (n=6) examined or regions of normal squamous esophageal epithelium within BE or EAC samples expressed GATA4 (Fig. 1). These data support a potential role for GATA4 in BE and EAC, either as a biomarker or, more interestingly, as a functional player in disease pathogenesis.

**Figure 1.**
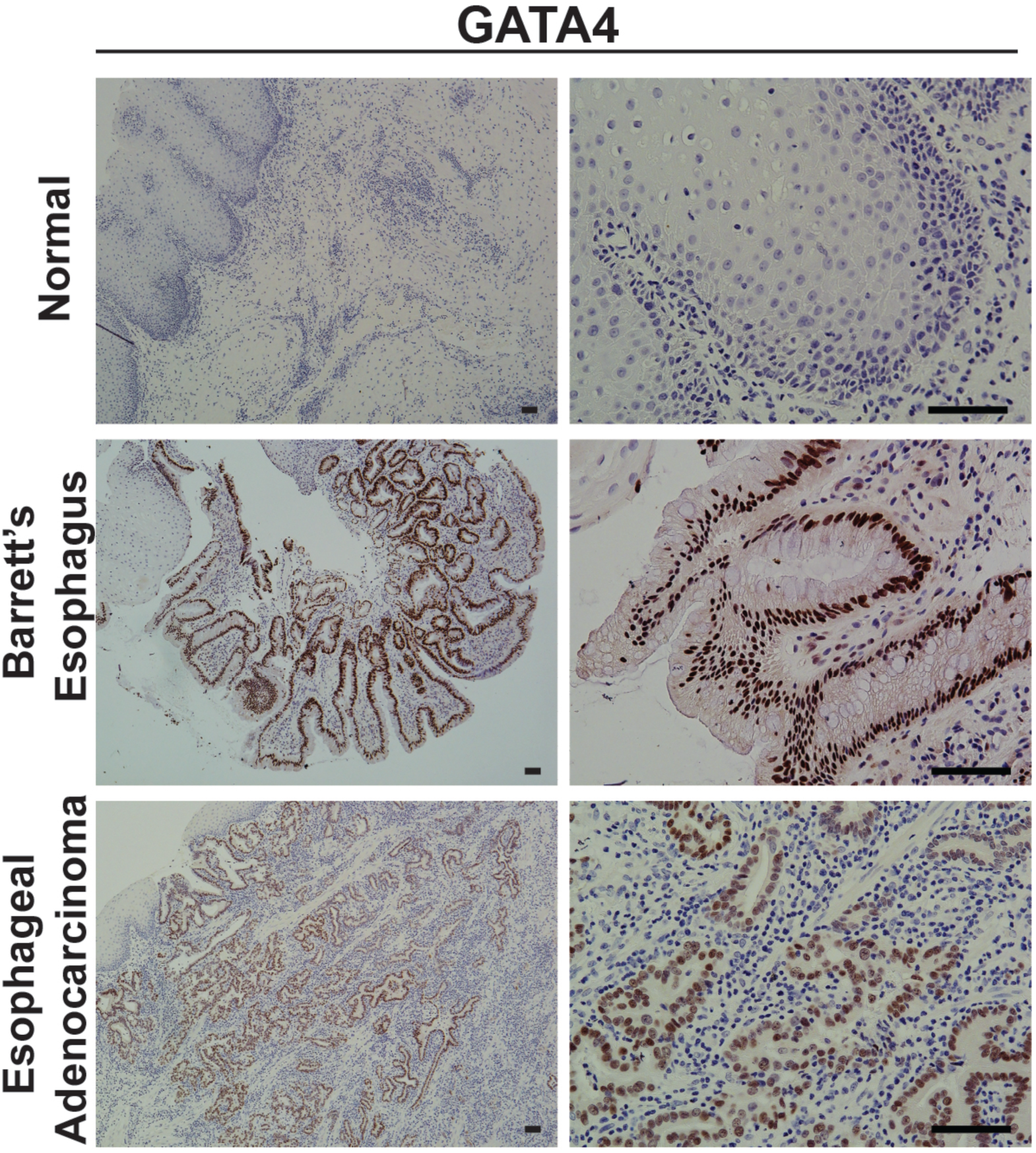
Human Barrett’s esophagus lesions and esophageal adenocarcinoma tumors aberrantly express GATA4. Tissue sections (5 µm) of de-identified human esophageal biopsy tissue without disease (n=5), with Barrett’s esophagus (n=10), or with esophageal adenocarcinoma (n=7) were obtained from the MCW Tissue bank and Department of Pathology. Immunohistochemistry was used to detect GATA4 protein (brown nuclear staining). No GATA4 protein was detected in normal tissue whereas all BE and EAC tissues analyzed expressed GATA4 protein in diseased epithelial cells. Sections were counterstained with hematoxylin. The presence of BE and/or EAC in tissue samples was verified by a board-certified GI pathologist. Scale bar, 100 µm

### Acid and bile induce GATA4 expression in human squamous cells

Previous studies have shown that treatment of human stratified squamous esophageal epithelial cells with acid and bile can induce expression of columnar cell markers including the intestinal transcription factor CDX2.[**30**] Therefore, to determine if acid and bile treatment would induce GATA4 in esophageal squamous epithelial cells, we treated an human esophageal squamous epithelial cell lines, NES-B10T, with acidified medium (pH 5.5) containing a mixture of bile salts for 48 hours and measured *GATA4* transcript levels by RT-PCR. We found that acid and bile treatment increased steady-state *GATA4* mRNA compared with control untreated cells (Fig. 2).

**Figure 2.**
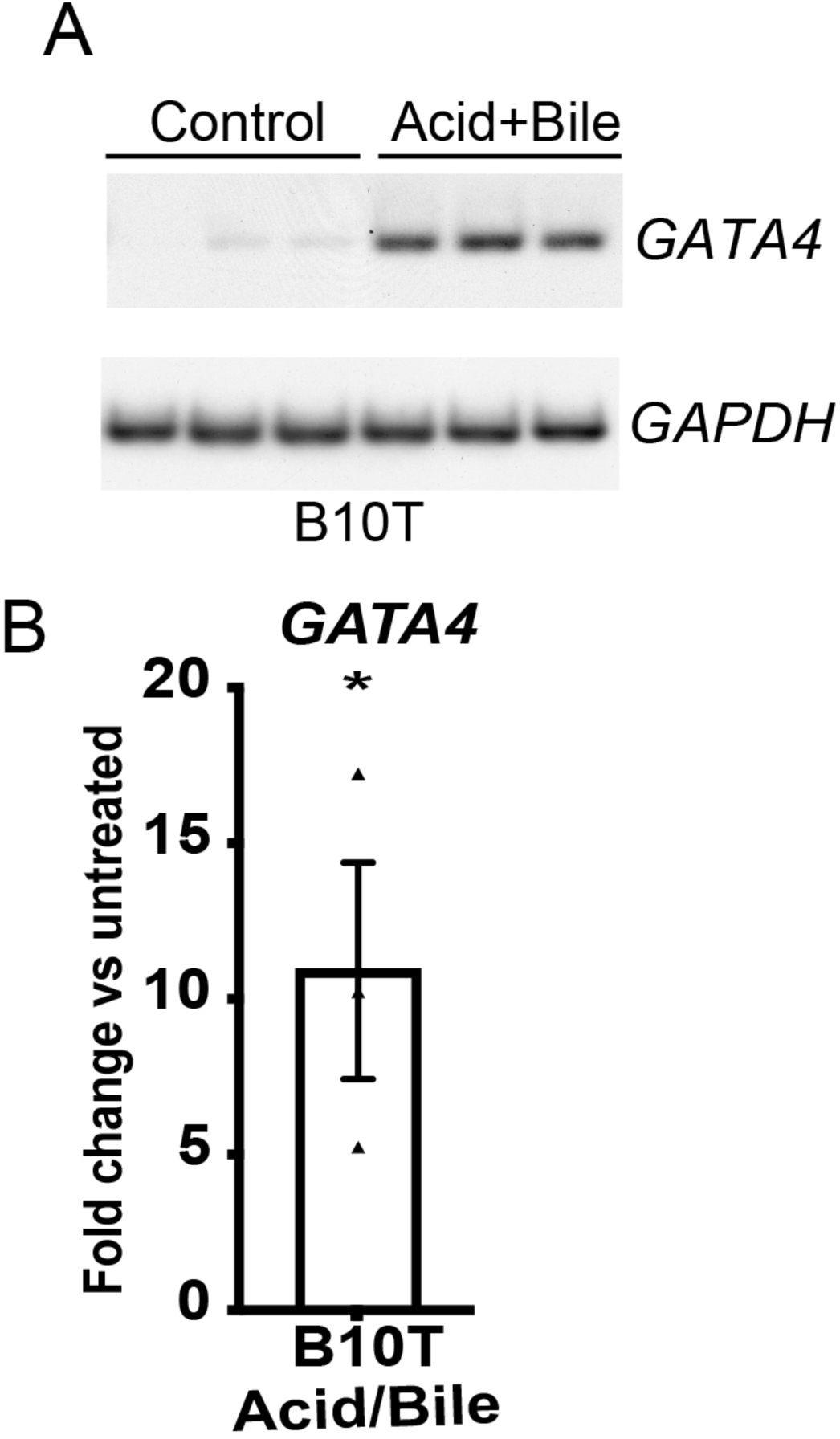
Acid and bile treatment of human esophageal cells in culture induces *GATA4* mRNA expression. The human esophageal squamous epithelial cell line NES-B10T was treated with acidified media (pH 5.5) containing a mixture of bile salts (20:3:15:3:6:1, final concentration 400 μM: glycocholic, taurocholic, glycochenodeoxycholic, taurochenodeoxycholic, glycodeoxycholic, and taurodeoxycholic acids) for 48 hours prior to RNA extraction and RT-PCR in the presence of [α-_32_P] dATP to detect the steady-state level of *GATA4* mRNA. Amplicons were separated by PAGE and visualized using autoradiography and phosphorimaging. (A) Representative image showing *GATA4* mRNA induced by acid and bile treatment of NES-B10T cells. *GAPDH* was used for normalization. (B) Phosphorimager quantification of data from three independent experiments demonstrated that *GATA4* mRNA was induced 10.9-fold in acid and bile treated NES-B10T cells (n=3; *P ≤ 0.05) compared with non-treated control cells.

### Ectopic GATA4 expression in NES-B10T cells reduced squamous cell marker mRNA expression and p63 protein

Our observations that acid and bile induced expression of GATA4, a columnar cell type transcription factor, in squamous epithelial cells and that GATA4 was abnormally present in human esophageal diseases suggested GATA4 as a candidate transcription factor in GERD-driven esophageal epithelial cell changes. Therefore, to explore the hypothesis that GATA4 plays a role in epithelial cell transformation related to BE, we used lentivirus to introduce a doxycycline-regulated GATA4 expression construct or a control construct lacking the GATA4 coding sequence (vector control) into NES-B3T and NES-B10T cells. Individual cell clones with stable integration of the expression construct (n=3 per cell line) or empty vector construct (n=1 per cell line) were treated with doxycycline (1 μg/ml) for 72 hours. We used qRT-PCR, immunoblot, and immunostaining to assess GATA4 mRNA and protein expression in each cell clone (Fig. 3). *GATA4* mRNA expression was detected in all cell clones containing the inducible expression construct and was undetectable in control cell clones (Fig. 3A). We also observed that higher levels of *GATA4* mRNA were generally present in NES-B10T cells transduced with the GATA4 expression construct compared with NES-B3T cells, with NES-B10T cell clones expressing an average of 5.3-fold more *GATA4* mRNA than NES-B3T cell clones (Fig. 3A).

**Figure 3.**
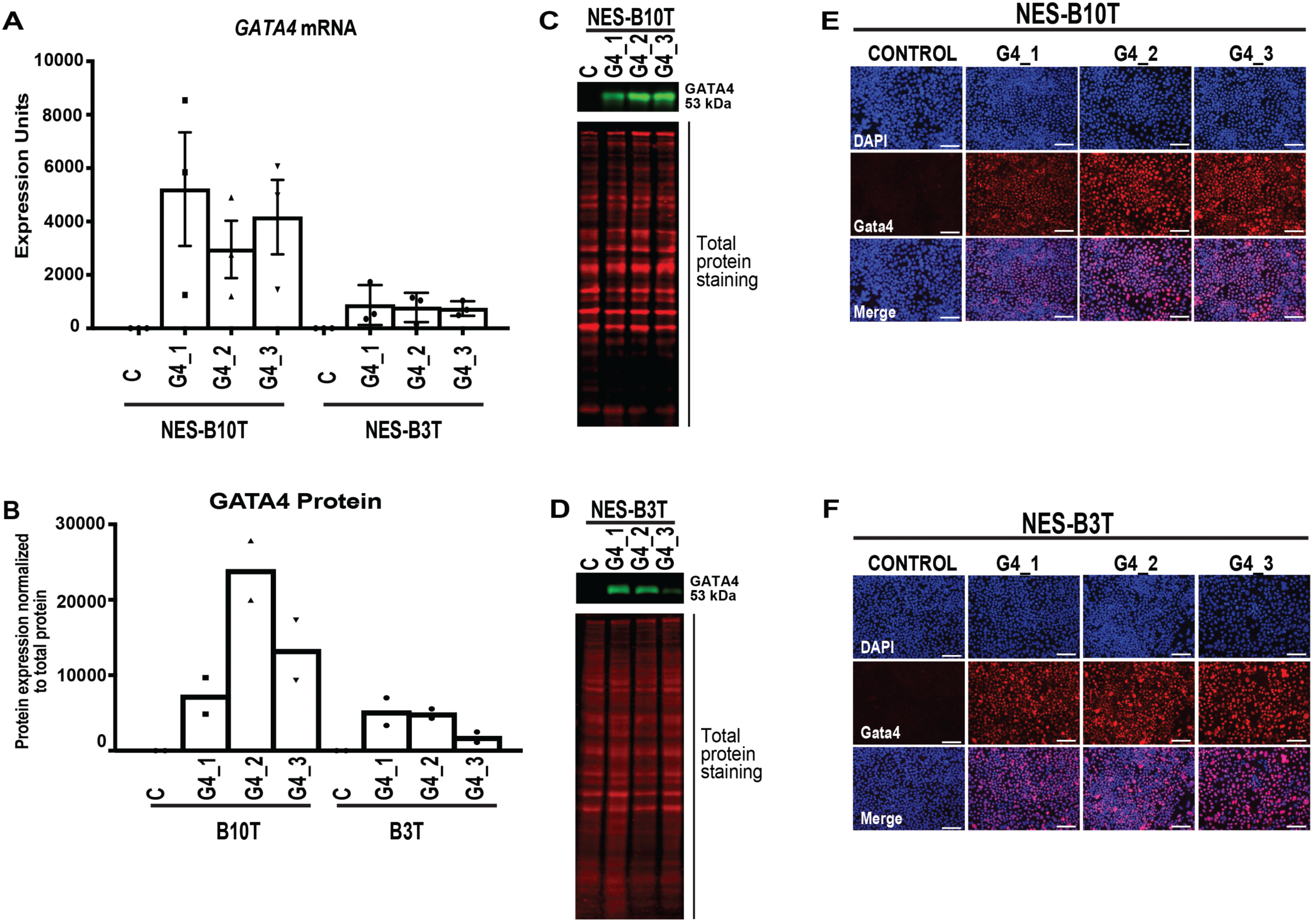
Generation of human esophageal squamous epithelial NES-B3T and NES-B10T cell clones with doxycycline-inducible expression of GATA4 protein. The human esophageal squamous epithelial cell lines NES-B10T and NES-B3T were infected with pInducer20 or pInducer20-GATA4 lentivirus (MOI 3), and clones were isolated after selection with G418. RNA and protein were collected from cells 72 hours post doxycycline (1 µg/ml) treatment. (A) qRT-PCR demonstrated *GATA4* mRNA induction in three pInducer20-GATA4 clones compared with one control clone per cell line. Data shown represent three independent induction experiments. (B-D) Quantitative infrared immunoblotting (LI-COR) was used to analyze GATA4 protein expression in nuclear extracts from control and GATA4 expressing B3T and B10T cell clones. Revert Total Protein Stain was used for normalization. Immunoblots were performed using nuclear extracts from two independent doxycycline induction experiments. Representative blots are shown in panels C and D. (E-F) Immunofluorescence staining of GATA4 protein expression (red nuclear staining) in three pInducer20-GATA4 clones compared with one control clone per cell line. DAPI (blue) indicates nuclei. IF detection for GATA4 protein was performed for one induction experiment to determine the distribution of GATA4 expressing cells. Scale bar, 100 µm.

To examine GATA4 protein expression in control and experimental cell clones, we used immunoblotting (Fig. 3B-D) and immunofluorescent protein staining (Fig. 3E-F). GATA4 protein was undetectable in control cell clones containing only the empty vector. In contrast, GATA4 protein was present in all six cell clones transduced with the GATA4 expression construct after doxycycline treatment. Similar to what was observed between cell lines at the mRNA level, we found that GATA4 protein levels, as measured by immunoblot, were higher in NES-B10T cells transduced with the GATA4 expression construct compared with NES-B3T cells similarly transduced, with NES-B10T cells expressing an average of 3.3-fold more GATA4 protein than B3T clones (Fig. 3B).

To determine the effect of GATA4 on expression of key markers of a squamous epithelial cell phenotype, we used qRT-PCR and immunoblotting to measure steady-state mRNA and/or protein levels of four well-studied and characteristic squamous epithelial cell marker genes, *p63, KRT5, KRT13*, and *KRT15*, in control and GATA4-induced NES-B10T and NES-B3T cell clones. We found that expression of GATA4 in NES-B10T cells decreased the abundance of all four mRNAs (Fig. 4A-D), although decreases in *p63* and *KRT5* in clone NES-B10T-G4-2 compared with control cells did not reach statistical significance (p=0.21 and p=0.06, respectively). We further examined how changes in mRNA affected the abundance of p63, KRT5, and KRT13 proteins. We observed that NES-B10T clones expressing GATA4 had reduced expression of p63 protein in all clones, but that KRT levels remained unchanged (Fig. 4 E-I).

**Figure 4.**
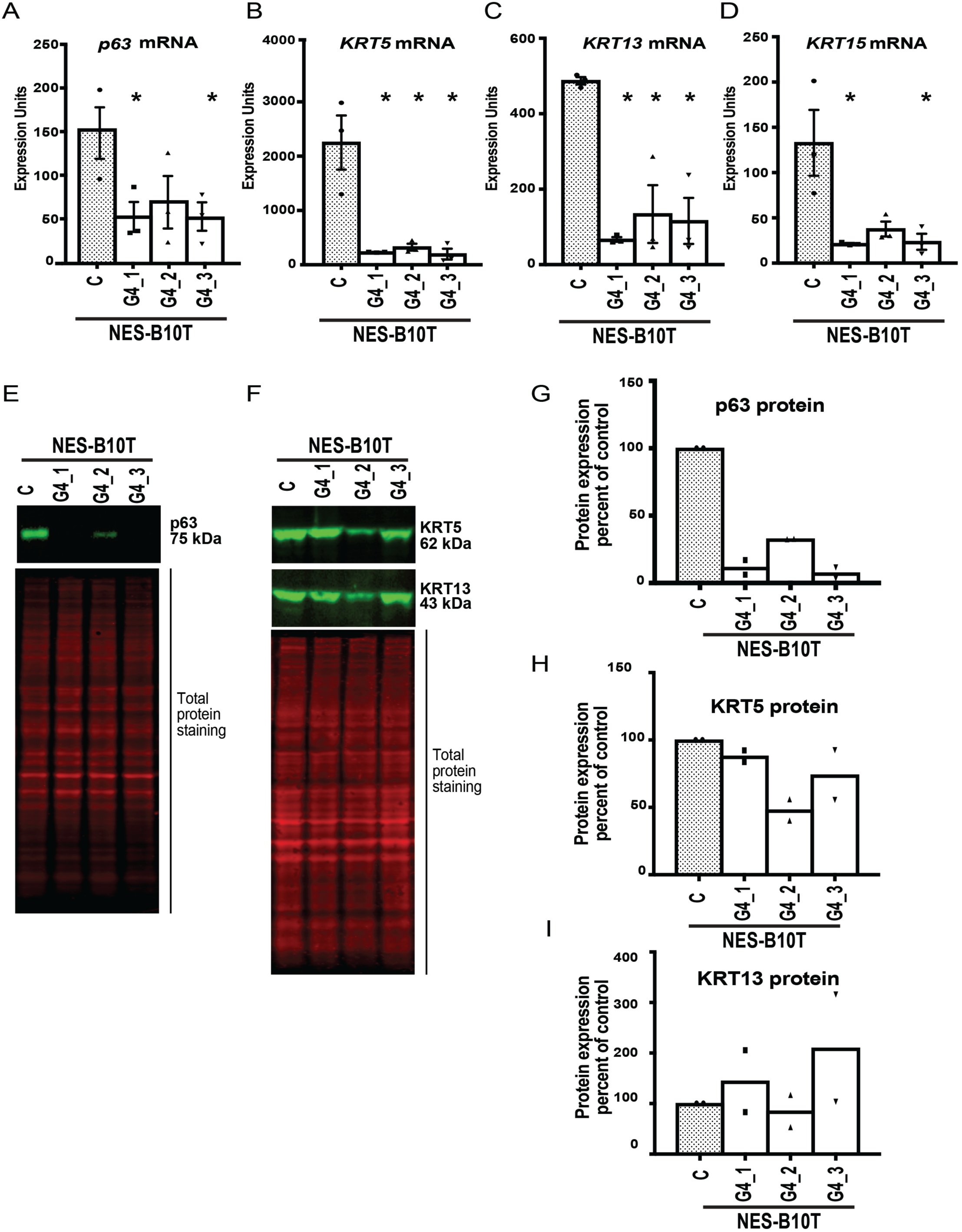
Ectopic expression of GATA4 in human squamous esophageal epithelial NES-B10T cells alters the expression of squamous cell marker genes. Expression of four squamous epithelial cell marker genes *(p63, KRT5, KRT13, KRT15*) were examined by qRT-PCR and/or immunoblot in three pInducer20-GATA4 NES-B10T cell clones (G4_1, G4_2, G4_3) and one pInducer20 NES-B10T control cell clone (C) after 72 hours of doxycycline treatment to induce GATA4. (A-D) qRT-PCR demonstrated reduced steady state levels of *p63, KRT5, KRT13*, and *KRT15* mRNA in NES-B10T cells expressing GATA4 compared with controls cells lacking GATA4 protein (n=3 experiments; *P ≤ 0.05). (E-I) Quantitative infrared immunoblotting (LI-COR) was used to analyze p63, KRT5, and KRT13 protein expression in control and GATA4 expressing NES-B10T cell clones. Revert Total Protein Stain was used for normalization. Immunoblots were performed using protein extracts from two independent doxycycline induction experiments. Representative blots are shown in panels E and F. Quantification of blots is shown in panels G-I with abundance of protein detected in GATA-expressing clones expressed as the percent relative to the non-expressing control cells.

We evaluated the same squamous epithelial cell marker gene expression in NES-B3T cell clones expressing GATA4. Overall, we found that expression of GATA4 in NES-B3T cells generally did not alter the abundance of any of the four marker mRNAs (Fig. 5A-D) or the abundance of p63, KRT5, or KRT13 proteins (FIG 5E-I). Only clone NES-B3T-G4-2 showed a statistically significant change in *p63* and *KRT5* mRNA levels (FIG 5A-B). This clone also had a somewhat reduced level of p63 protein compared with control cells (Fig. 5E,G). We attributed the different effects of GATA4 in these two independently derived cell lines to the different levels of GATA4 induced in the cell lines. We also observed a difference in the baseline expression of p63 protein between NES-B10T and NES-B3T control cell clones. After normalizing to total protein, we found that NES-B3T control cells contained approximately 10-fold more p63 protein compared with NES-B10T control cells. The increased baseline level of p63 in NES-B3T cells could make NES-B3T cells more resistant to transformation by GATA4.

**Figure 5.**
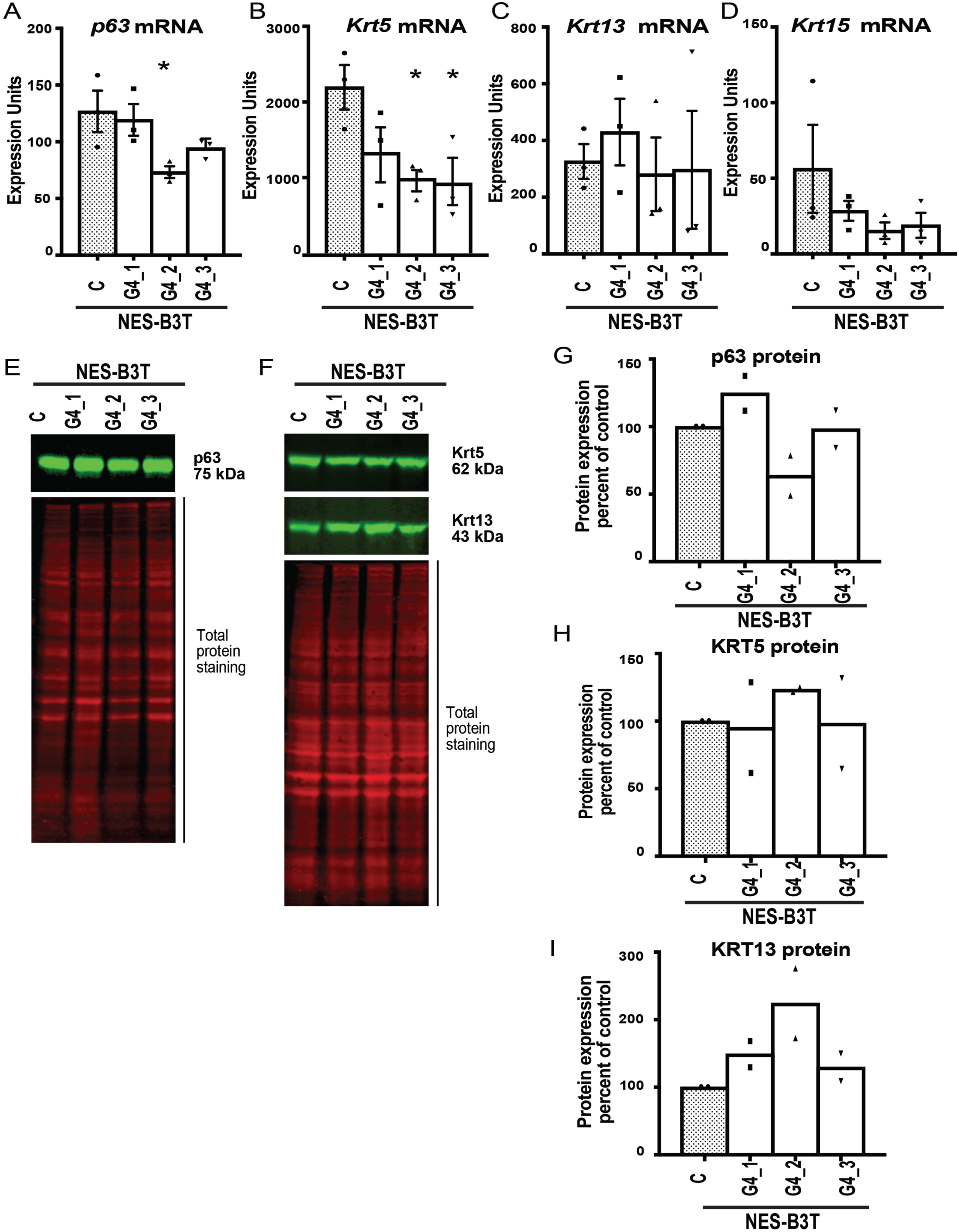
Effects of ectopic expression of GATA4 on expression of squamous epithelial cell marker genes in human squamous esophageal epithelial NES-B3T cells were less pronounced than those observed in GATA4-expressing NES-B10T cells. Expression of four squamous epithelial cell marker genes *(p63, KRT5, KRT13, KRT15*) were examined by qRT-PCR and/or immunoblot in three pInducer20-GATA4 NES-B3T cell clones (G4_1, G4_2, G4_3) and one pInducer20 NES-B3T control cell clone (C) after 72 hours of doxycycline treatment to induce GATA4. (A-D) qRT-PCR demonstrated a trend for reduction in the steady state levels of *p63, KRT5*, and *KRT15*, but not *KRT13*, in NES-B3T cells expressing GATAA4 compared with controls cells lacking GATA4 protein. Statistically significant reductions were observed for only one cell clone, G4_2, for *p63* expression and two clones, G4_2 and G4_3, for *KRT5* expression (n=3 experiments; *P ≤ 0.05). (E-I) Quantitative infrared immunoblotting (LI-COR) was used to analyze p63, KRT5, and KRT13 protein expression in control and GATA4 expressing NES-B3T cell clones. Revert Total Protein Stain was used for normalization. Immunoblots were performed using extracts from two independent doxycycline induction experiments. Representative blots are shown in panels E and F. Quantification of blots is shown in panels G-I with abundance of protein detected in GATA-expressing clones expressed as the percent relative to the non-expressing control cells.

Observing that squamous type keratin gene expression was altered in GATA4 expressing esophageal epithelial cells, we examined mRNA expression of a typical columnar cytokeratin, *KRT8*, in control and GATA4-expressing NES-B3T and NES-B10T cell clones. We found that control NES-B3T and NES-B10T cell clones expressed low baseline levels of *KRT8* and that GATA4 expression increased *KRT8* expression in all clones by about 4-fold (Fig. 6). Compared with the effects on stratified squamous epithelial cell marker gene expression, induction of this columnar epithelial cell marker was more uniform across the two cell lines, even given variations in GATA4 levels across clones. Examination of mRNA expression of the other columnar markers *CDX2, Villin*, and *KRT20* by qRT-PCR showed none to be induced in these GATA4 expressing cells (data not shown).

**Figure 6.**
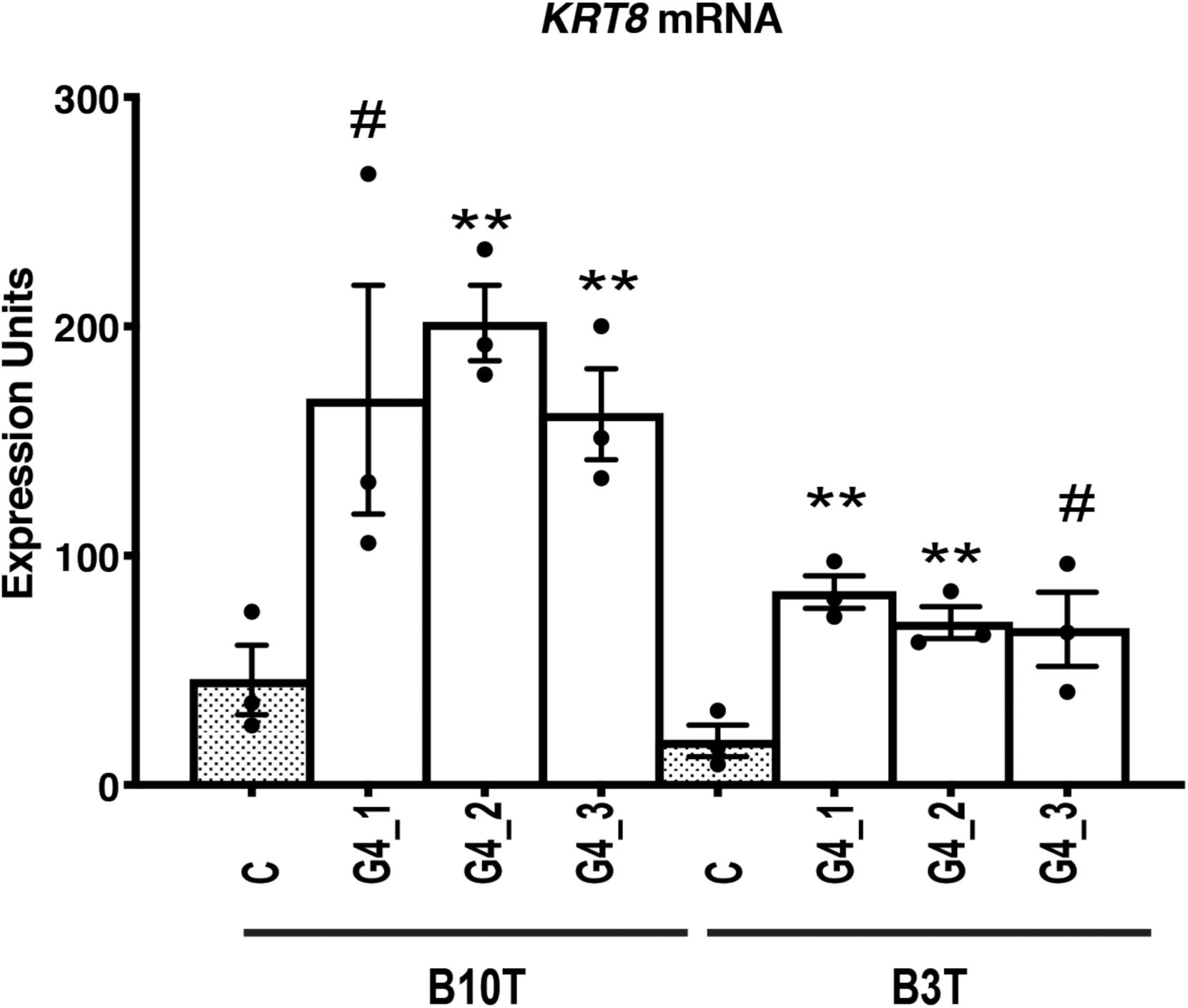
The columnar epithelial cell cytokeratin *KRT8* was induced in human esophageal epithelial NES-B3T and NES-B10T cells expressing GATA4 protein. Expression of the steady-state level of *KRT8* mRNA was examined by qRT-PCR in three pInducer20-GATA4 cell clones (G4_1, G4_2, G4_3) and one pInducer20 control cell clone (C) per cell line after 72 hours of doxycycline treatment to induce GATA4. qRT-PCR demonstrated up-regulation of *KRT8* mRNA in NES-B3T and NES-B10T cells expressing GATA4 compared with controls cells lacking GATA4 protein (n=3 experiments; **P ≤ 0.01; _#_P=0.0788, NES-B10T clone G4_1 and P=0.0503, NES-B3T clone G4_3).

### GATA4 binds to promoters of squamous epithelial cell marker genes

Our lab’s previous studies of GATA4 function at the jejunal-ileal intestinal boundary show that GATA4 can act to either activate or repress gene transcription.[**31**,**32**] In the jejunum, GATA4 occupies GATA consensus binding sites in chromatin to activate expression of genes typically expressed in the jejunum and to repress expression of genes typically expressed solely in the ileum. This work, along with studies of GATA4 function in tissues like the heart, sets a precedent for GATA4 to act as a transcriptional repressor. Therefore, we sought to determine if GATA4 occupies the promoters of the squamous epithelial cell marker genes that we found with repressed expression in NES-B10T GATA4-expressing cells—*p63, KRT5, KRT13*, and *KRT15*. Because of a lack of a well-validated GATA4 antibody for chromatin immunoprecipitation, we turned to a mouse model, *Gata4*_*flbio/flbio*_::*ROSA26*_*BirA/BirA*_, in which endogenous GATA4 protein is biotinylated (GATA4-BIO) allowing for precipitation of GATA4-bound chromatin complexes by streptavidin coated beads.[**33–35**] We chose to use mouse hindstomach because these columnar cells normally express GATA4 and lack expression of *p63, Krt5, Krt13*, and *Krt15* (Fig. 7A). In contrast, mouse forestomach squamous epithelial cells, akin to human esophageal cells, lack GATA4 and express these squamous marker genes (Fig. 7A). We performed bio-ChIP PCR using sonicated chromatin isolated from columnar hindstomach epithelial cells of *Gata4*_*flbio/flbio*_::*ROSA26*_*BirA/BirA*_ mice (biotinylated GATA4, GATA4-BIO) and *ROSA26*_*BirA/BirA*_ mice (non-biotinylated GATA4, GATA4-WT) (n=4 animals of each genotype). To identify relevant GATA4 binding sequences proximal to each gene’s transcriptional start site (TSS), we looked for evolutionarily conserved GATA consensus sequences in the promoters of these genes. We aligned genomic sequences containing 1KB upstream of each gene’s TSS from mouse, chimpanzee, and human and found one evolutionarily conserved GATA site in each of the *p63, KRT5*, and *KRT15* promoters and no conserved site in the *KRT13* promoter (Fig. 7B). We designed primers flanking the conserved GATA binding sites for PCR from isolated chromatin. For a negative control, we used primers to assay GATA4 occupancy at a region in the ubiquitously-expressed *Hprt1* gene that lacks GATA4 binding sites and that we have validated as a negative GATA4 ChIP control in other published studies.[**32**,**36**,**37**] GATA4 enrichment at the putative binding sites in squamous cell gene promoters in hindstomach chromatin from GATA4-BIO animals compared with that of control GATA4-WT was measured and expressed as a percentage of input (Fig. 7C). We observed amplification of the binding regions in the promoters of *p63, Krt5*, and *Krt15* only in chromatin isolated from GATA4-BIO animals and not from chromatin isolated from control animals (Fig. 7C). As expected, no enrichment was observed for the negative control *Hprt1* (Fig. 7C). These data suggest that GATA4 binds to sites in the *p63, KRT5*, and *KRT15* promoters in columnar epithelial cells to repress expression of these genes.

**Figure 7.**
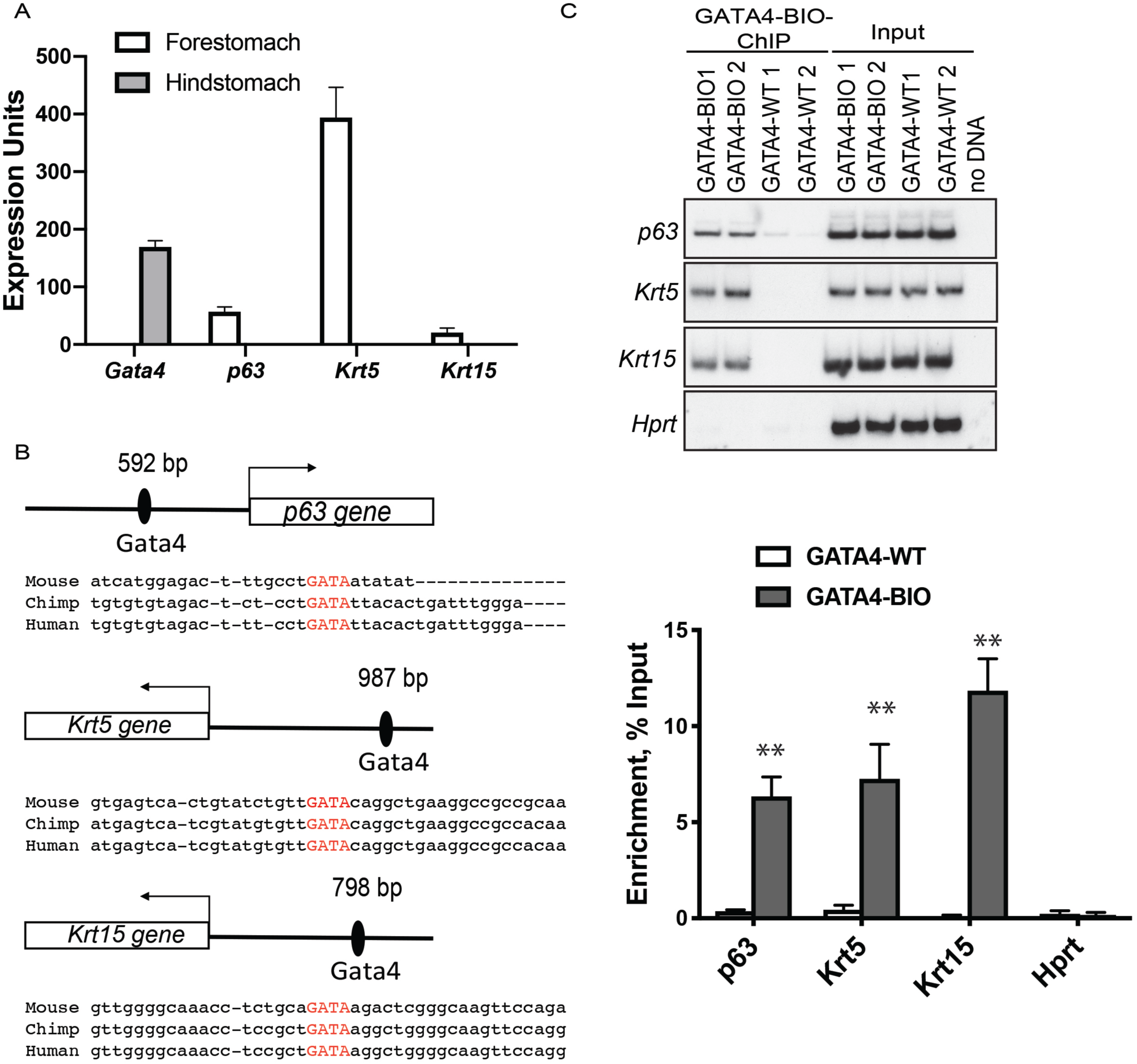
GATA4 occupies the proximal promoters of squamous epithelial cell marker genes suggesting that GATA4 can repress expression of these genes. (A) To validate mouse hindstomach epithelial cells as a model system to examine GATA4 binding to regulatory regions of genes marking stratified squamous epithelium to repress their expression, qRT-PCR was used to examine the expression profiles of *Gata4, p63, Krt5*, and *Krt15* genes in mouse columnar, glandular hindstomach and stratified squamous forestomach epithelial cells. As expected, *Gata4* transcript was detected only in columnar hindstomach epithelium whereas *p63, Krt5*, and *Krt15* transcripts were detected only in stratified squamous forestomach. (B) Diagrammatic representation of GATA consensus binding sites identified as evolutionarily conserved among human, mouse, and chimpanzee in the *p63, Krt5*, and *Krt15* genes. (C) GATA4 Bio-ChIP–PCR showed enriched amplification of regions containing predicted evolutionarily conserved GATA binding sites in the *p63, Krt5* and *Krt15* genes in cells expressing biotinylated GATA4 compared with control cells lacking biotinylated GATA4. No amplification enrichment was observed for a gene lacking a GATA4 binding site *(Hprt*). Autoradiographic band intensity was measured using a Storm820 Phosphor Imager and ImageQuant software. epresentative autoradiographic data for two of four animals per genotype assayed shown. Amplification enrichment per sample was normalized to input (n = 4 GATA4-BIO and 4 GATA4-WT). Error bars show SEM, \*\**P* ≤ 0.01.

## DISCUSSION

Our observation that human esophageal squamous epithelial cells respond to acid and bile by inducing *GATA4* mRNA expression suggests that cells residing within the esophagus in vivo would similarly have the capacity to ectopically induce GATA4 expression in the presence of acid and bile, providing a possible link between chronic GERD and GATA4 expressing metaplastic cells. Although it currently remains unclear whether or not resident esophageal epithelial cells contribute to the origin of BE, these data support the idea that an acid and bile-rich reflux environment could support and stabilize abnormal GATA4-expressing cells.

It is clear from our work that individual human esophageal cell lines react uniquely to ectopic expression of GATA4. We found that NES-B10T cells accommodated a higher level of GATA4 mRNA and protein expression compared with NES-B3T cells. This result is unlikely due simply to positional effects related to lentiviral integration, as three independent cell clones from each line behaved similarly. It more likely reflects inherent differences between these cell lines, and, more globally, may reflect differences between the two individuals from which these lines were derived. Moreover, this result suggests post-transcriptional and post-translational regulation of GATA4 levels given that the expression construct contains only the GATA4 coding sequence and, therefore, lacks endogenous GATA4 regulatory elements. A recent study demonstrates that GATA4 protein is subject to selective autophagic degradation, suggesting that there could be differences in selective autophagy of GATA4 between these two cell lines.[**38**]

Our data further suggest a dose dependent effect of GATA4 on human squamous esophageal epithelial cells. In NES-B10T GATA4 cell clones, which expressed about three times more GATA4 protein than NES-B3T GATA4 cell clones, we saw changes in expression of the typical stratified squamous cell marker transcripts *p63, KRT5, KRT13*, and *KRT15*. We further observed a strong downregulation of p63 protein in NES-B10T cells. KRT5 and KRT13 protein levels were not changed in this acute three-day GATA4 induction treatment, and this likely reflects the longer half-life of these cytoskeletal proteins. In general, cytokeratins are long-lived cellular proteins with the average half-life of each spanning around 100 hrs, which is necessary for their function as cytoskeletal building blocks.[**39**] In contrast, the transcription factor p63, is short-lived with a half-life of 10-40 hrs.[**40**]

Supporting the idea that NES-B10T and NES-B3T harbor important differences that affect their ability to be transformed by GATA4 was our finding that baseline p63 levels differ significantly between these cell lines with NES-B3T cells expressing approximately 10-fold more p63 than NES-B10T cells. This may be attributable to genetic differences between the donors, or it could reflect differences in physiological or environmental exposures between the individuals from which the cells were derived. One consequence of higher levels of p63 could be resistance to transformation by GATA4 as we observed that NES-B3T cells did not sustain high GATA4 levels and consequentially showed more subtle effects in stratified cells marker expression compared with NES-B10T cells. Moreover, a higher level of p63 in NES-B3T cells could make these cells more resistant to GATA4 induction given that p63 has been implicated in regulating autophagy and GATA4 is a protein targeted for degradation via autophagy.[**38**,**41**]

Although we observed alterations in expression of multiple squamous cell marker genes, we only observed up-regulation of one of the columnar cell marker gene we analyzed, *KRT8*. It was interesting that the effect on *KRT8* was more uniform across the NES-B3T and NES-B10T cell lines even with the different levels of GATA4. It is possible that longer exposure to GATA4, beyond the acute three-day window examined in this study, would result in further transformation toward a columnar cell fate. In fact, a study in which NES-B10T cells were treated with acid and bile for 5 min/day for 30 weeks showed that expression of markers of columnar epithelium increased progressively over time.[**13**] Future experiments could test this by inducing GATA4 for longer time periods.

Our data suggest that GATA4 exerts its effects in esophageal epithelial cells by directly binding to GATA consensus binding sites in the promoters of the *p63, KRT5*, and *KRT15* genes to repress expression of these squamous marker transcripts. Although we did not identify an evolutionarily conserved binding site in the *KRT13* gene promoter, this does not rule out the presence of GATA4 regulatory sites beyond the promoter, perhaps in an enhancer. GATA4 is known to activate or repress gene expression. Our laboratory’s previous work examining GATA4’s role in defining the jejunal-ileal epithelial boundary illustrates well how GATA4 acts on one set of genes in a positive manner to promote a particular gene expression program and cellular identity while simultaneously acting on another set of genes in a negative manner to repress another undesired gene expression program and its concomitant cellular identity.[**31**,**32**] Although we were unable to directly assess GATA4 binding to these promoters in NES-B10T cells because of the lack of a reliable GATA4 ChIP antibody, the evolutionary conservation of these binding sites supports the premise that GATA4 directly represses expression of these genes when expressed in human esophageal cells. It is possible that the differential repressive effects we noted on expression of squamous epithelial cell genes between the two cell lines reflects differences in expression of essential GATA4 co-factors cooperating with GATA4 to repress squamous gene expression. Taken together, our data suggest a mechanism whereby GATA4 expression in abnormal esophageal cells, possibly induced by reflux conditions, could support a columnar metaplastic cell identity program by repressing expression of key genes required for stratified cell identity.

## METHODS

### Tissue immunohistochemistry

Sections (5 µm) of formalin-fixed, paraffin embedded de-identified human tissue patient biopsy samples were obtained through the MCW Tissue Bank and Department of Pathology and stained with hematoxylin and eosin. The presence of Barrett’s esophagus or esophageal adenocarcinoma in tissues was verified by a board-certified GI pathologist. Standard immunohistochemical procedures were followed.**[32**,**45]** Briefly, citric acid antigen retrieval was performed prior to immunohistochemistry for GATA4 (R&D Systems, AF2606). Staining was visualized using R.T.U. Vectastain Elite ABC reagent and a Metal Enhanced DAB substrate kit. Micrographs were captured using a Nikon Eclipse TE300 fluorescent microscope and SPOT RT3 camera. Images were assembled into figures using Adobe Photoshop and Illustrator. Images from control and experimental samples were processed identically.

### Cell culture

The NES-B3T and NES-B10T telomerase-immortalized human squamous esophageal cell lines were kindly provided by Dr. Rhonda Souza.[**42**,**43**] Both cell lines were established from biopsy samples of individual patients with Barrett’s esophagus, but not from the diseased region. For expansion and maintenance, cells were co-cultured with mitomycin-C inactivated mouse embryonic fibroblast feeder cells in DMEM/F12 (3:1) medium containing hydrocortisone (0.4 μg/ml), EGF (20 ng/ml), transferrin (5 μg/ml), insulin (5 μg/ml), cholera toxin 10 ng/ml, triiodothyronine (20 pM), adenine (31 μg/ml), cosmic calf serum (1%), penicillin/streptomycin (100 U/ml each). Cells were cultured at 37°C in a humidified 5% CO_2_ incubator. For experiments, cells were cultured on collagen IV coated plates in the same medium.

### Acid and bile treatment of cells

Cells (70% confluency) were treated with acidified medium (pH 5.5) containing bile salts (20:3:15:3:6:1, final concentration 400 μM: glycocholic, taurocholic, glycochenodeoxycholic, taurochenodeoxycholic, glycodeoxycholic, and taurodeoxycholic acids) for 48 hours as previously described.[**30**] The pH of the medium was adjusted to pH 5.5 immediately before treatment using 1M hydrochloric acid. After 48 hours, cells were flash frozen in RLT buffer containing β-mercaptoethanol for RNA extraction.

### Lentiviral GATA4 vector transduction and cloning

NES-B3T and NES-B10T cells were infected with pInducer20 or pInducer20-GATA4 lentivirus (MOI 3) prepared by the viral core of the Medical College of Wisconsin. Plasmids were generously provided by Dr. Stephen Elledge (Harvard Medical School).[**38**] Infected cells were maintained in culture medium supplemented with Tet system approved FBS (5%). Cells were selected with G418 (350 μg/ml, B3T; 500 μg/ml, B10T) beginning 24 hours after infection. The G418 dose was established as double the concentration required to effectively kill non-infected cells. To obtain individual clones, a limiting dilution cloning method was applied, and cells were plated onto collagen IV coated 96-well plates. For experiments, doxycycline (1 μg/ml) was applied to all cell cultures (control and GATA4-transduced) for 72 hours. Media was replaced after 48 hours to replenish doxycycline. Cells were treated with trypsin to harvest for RNA and protein extraction or fixed with 4% PFA for immunofluorescence experiments.

### Reverse-Transcription PCR (RT-PCR)

Total RNA was isolated using RNeasy Mini Kit. To remove genomic DNA, total RNA was treated with ezDNase. For quantitative reverse-transcription polymerase chain reaction (qRT-PCR), cDNA synthesized using MMLV and random hexamer primers was amplified using TaqMan Gene Expression Mastermix and TaqMan gene expression assays (*GATA4*, HS00171403_m1; *KRT5*, Hs00361185_m1; *KRT8*, Hs01595539_m1; *KRT13*, Hs00357961_g1; *KRT15*, Hs00951967_m1; *KRT20*, Hs00300643_m1; *p63*, Hs00978340_m1; *CDX2*, Hs01078080_m1; *VILLIN*, Hs01031724_m1). Data were normalized to expression of glyceraldehyde-3-phosphate dehydrogenase (*GAPDH*, 4352665). For each target, expression units were calculated using the formula [2_(-δCq)_]x1000.[**44**] For radioactive RT-PCR with [α-_32_P] deoxyadenosine triphosphate, cDNA was generated from ezDNase treated RNA using a Superscript VILO cDNA synthesis kit. Primers used were *GATA4* (5’-ttctggggagagtgtaagtggacag-3’, 5’-ctttttgcctcctggacaaaagact-3’) and *GAPDH* (5’-gacagtcagccgcatcttct-3’, 5’-ttaaaagcagccctggtgac-3’). Radioactive amplicons were separated by 4% PAGE, and gels were dried. Expression was detected and quantified using Storm80 Phosphor Imager (Amersham Biosciences).

### Immunocytochemistry

Standard immunocytochemistry methods were used.**[45]** Briefly, cells were fixed with 4% PFA/1X PBS, permeabilized with 0.5% Triton X-100/1X PBS, blocked with 3% BSA/1X PBS, and incubated with antibodies (GATA4, R&D Systems, AF2606; Alexa-Fluor 594 Donkey anti-goat, Thermo Fisher) in 1% BSA/1X PBS. DAPI was used to visualize nuclei. Images were taken with Nikon Eclipse TE300 fluorescent microscope equipped with SPOT RT3 camera with 20X objective. Images were assembled into figures using Adobe Photoshop and Illustrator. Images from control and experimental samples were processed identically.

### Immunoblot analysis

Standard immunoblot methodology was followed.**[32**,**45]** Nuclear extracts were prepared using the NE-PER Nuclear and Cytoplasmic Extraction Reagents and HALT protease inhibitor cocktail. Benzonase (0.5 U/μl) was used to increase protein yield of transcription factors and quality of nuclear extracts. Nuclear extracts (5 μg) or cytoplasmic extracts (20 μg) were separated using Nu-PAGE Bis-Tris 4%-12% gradient gels and transferred to an Immobilon-FL polyvinylidene difluoride membrane. Total protein was detected by Revert Total Protein Stain. Blots were blocked with Odyssey blocking buffer for 1 hour at room temperature. Antibody (GATA4 D3A3M, Cell Signaling Technology, #36966; KRT5, Covance, #PRB-160P; KRT13, Abcam, #92551; p63 D2K8X, Cell Signaling Technology, #13109) was added to blocking buffer containing 0.1% Tween and rotated at 4°C overnight. Membranes were washed and exposed to secondary antibody (IRDye 800CW, donkey anti-rabbit, LICOR) for 1 hour at room temperature. Blots were visualized using an Odyssey Infrared Imaging System (LI-COR) and quantitated using Odyssey software using normalization to REVERT total protein stain. If required, membranes were stripped with 2% SDS, 62.5 mM Tris buffer, pH 6.8, containing 0.8% β-mercaptoethanol at 50° C for 45 min. Stripped membranes were washed extensively under running tap water for 2 hr. Total protein was requantified using REVERT total protein stain, and proteins of interest were detected as described above.

### Animals

*Gata4*_*flbio/flbio*_(*Gata4*_*tm3*.*1Wtp*_) and *ROSA26*_*BirA*_*(Gt*(ROSA)*26Sor*_*Tm1[birA]Mejr*_) mouse lines were used to generate *Gata4*_*flbio/flbio*_::*ROSA26*_*BirA/BirA*_ (GATA4-BIO, biotinylated GATA4) and *ROSA26*_*BirA/BirA*_ (GATA4-WT, non-biotinylated GATA4) mice for biotin-mediated chromatin immunoprecipitation PCR (bio-ChIP-PCR)[**32–35**]. Primers for PCR genotyping of ear punch DNA were Gata4 Exon 7F/R, 5’-cagtgctgtctgctctgaagctgt-3’, 5’-ccaaggtgggcttctctgtaagaac-3’; BirAJax 14/15 5’-ttcagacactgcgtgact-3’, 5’-ggctccaatgactatttgc-3’; and BirAJax 16/17 5’-gtgtaactgtggacagaggag-3’, 5’-gaacttgatgtgtagaccagg-3’.[**32**] The Institutional Animal Care and Use Committee of the Medical College of Wisconsin approved all animal procedures. All experiments were performed in accordance with relevant guidelines and regulations.

### Bio-ChIP-PCR

Hindstomach epithelial cells from 2-4-month-old GATA4-BIO or GATA4-WT mice (n = 4 animals per genotype) were obtained as previously described with minor changes.[**32**] Briefly, mouse hindstomach epithelia were separated into single cells by vortexing dissected, washed hindstomach tissue for 30 minutes at 4°C in a balanced sodium salt solution (BSS) buffer with EDTA (1.5 mmol/L KCl, 96 mmol/L NaCl, 27 mmol/L Na_3_C_6_H_5_O_7_, 8 mmol/L KH_2_PO_4_, 5.6 mmol/L Na_2_HPO_4_, and 15 mmol/L EDTA, 200 μmol/L phenylmethylsulfonyl fluoride). Following removal of mesenchymal and muscle tissues, epithelial cell isolates were pelleted by centrifuging at 2000 rpm for 8 minutes at 4°C. Cells were washed twice with 1X PBS containing 200 μmol/L phenylmethylsulfonyl fluoride. Bio-ChIP was performed per previously published protocols.**[32**,**34**,**37]** Briefly, cells were fixed with 1% formaldehyde for 10 minutes, quenched with glycine (final concentration 125 mM), and flash frozen. Cells were lysed and sonicated in 2% SDS buffer using Bioruptor Pico for 6 total cycles, 30 sec sonication/30 sec rest for each cycle. GATA4 bound chromatin was isolated using magnetic streptavidin beads. After washes and proteinase K/ribonuclease A treatment, chromatin was purified by phenol/chloroform extraction and ethanol precipitation. Chromatin shearing size was determined by Agilent 2200 TapeStation with high sensitivity screen tape. Concentration was measured by Qubit™ dsDNA HS Assay Kit. Evolutionarily conserved GATA4 binding sites were identified by aligning 1KB upstream of each gene’s transcriptional start site among mouse, human, and chimpanzee. GATA4 occupancy of evolutionary conserved GATA binding sites in the upstream regulator regions the *p63, Krt5* and *Krt15* genes was validated by radioactive PCR with [α-_32_P] deoxyadenosine triphosphate and gene specific primers (*p63*, 5’-gcctacattcagaaaggaaacaaattc-3’, 5’-gcctgtcaatggggaaaaataaagt-3’; *Krt5*, 5’-tattccaccagggaagacgtgagt-3’, 5’-tttgggccagagatagaggaaacac-3’; *Krt15*, 5’-aagtctgattgatcaccctgtcacc-3’, 5’-tttctggaacttgcccgagtcttat-3’). As a negative control, GATA4 enrichment was assessed using primers for an exon in the *Hprt* gene (5’-agcgcaagttgaatgtgc-3’, 5’-agcgacaatgtaccagag-3’), which we had previously validated as spanning a region without GATA4 binding.[**32**,**36**,**37**] Amplicons were separated using 4% PAGE and gels were dried. Enrichment was quantified using Storm820 Phosphor Imager (Amersham Biosciences) as previously described.[**32**] Percentage enrichment was calculated using the value for the input sample to normalize.

### Statistical analysis

Data were analyzed using an unpaired Student’s t-test when appropriate. A P-value of less than or equal to 0.05 was considered as statistically significant for observed changes. Standard error of the mean (SEM) is shown on graphs when appropriate.

## Acknowledgments

We thank Ugochukwu Ihenacho (Medical College of Wisconsin) for technical assistance. We thank Dr. Stephen Elledge (Harvard Medical School) for pInducer20 and pInducer20-GATA4 plasmids. This work was supported by the National Institutes of Health (NIH), National Institute of Diabetes and Digestive and Kidney Diseases (DK111822, MAB) and the Medical College of Wisconsin Cancer Center (MAB).

---

**Supplemental Figure 1.** Full immunoblots from which figure panels were derived. (A) Immunoblot corresponding to Figure 3C; (B) Immunoblot corresponding to Figure 3D; (C) Immunoblot corresponding to Figures 4E and 5E; (D-E) Immunoblots corresponding to Figures 4F and 5F.

## Data Availability Statement

No datasets were generated or analyzed during the current study.

