## Supplementary figures and images for "GATA4, expressed in Barrett’s esophagus and esophageal adenocarcinoma, can block squamous epithelial cell gene expression in human esophageal cells"

### Supplemental Figure 1

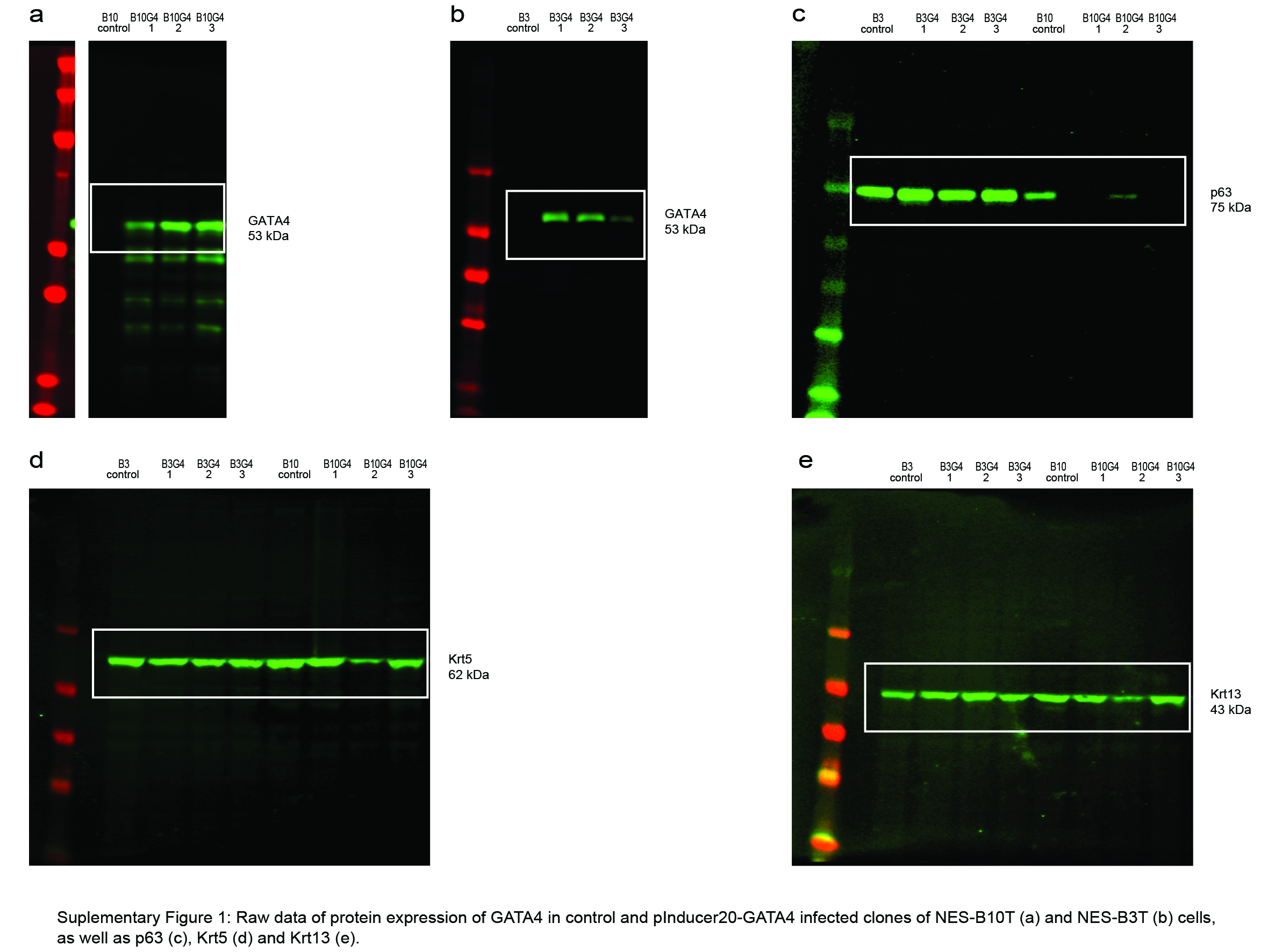
